# Supplementation with effective microorganisms in earthen ponds affects common carp growth but not overall microbial communities

**DOI:** 10.1101/2025.07.07.663436

**Authors:** Michalina Jakimowicz, Katarzyna Sidorczuk, David Huyben, Falk Hildebrand, Łukasz Napora-Rutkowski, Piotr Hajduk, Marek Sztuka, Magda Mielczarek, Dawid Słomian, Urszula Szulc, Laura Jarosz, Joanna Szyda

## Abstract

Probiotics are increasingly explored in aquaculture to enhance fish health and growth without leaving harmful residues. However, their efficacy in real-world pond environments remains poorly understood. Here, we conducted a 103-day field experiment to assess the effects of two effective microorganisms commercial products supplementations on microbial communities and growth performance of common carp (*Cyprinus carpio*). Effective microorganisms were added both to feed and directly to pond water. Microbial diversity was analysed using 16S rRNA and whole-genome shotgun sequencing across three environments – water (three time points), sediment (two time points) and fish intestine (one time point) – from 25 experimental ponds. Bioinformatics processing involved QIIME2 and MG-TK pipeline with taxonomic classification based on the SILVA database. The results showed that although supplemented bacterial families did not establish significantly in pond environments, fish exposed to specific effective microorganisms treatments exhibited improved growth metrics. These findings suggest that effective microorganisms can enhance carp growth in aquaculture without significantly altering resident microbial communities, offering a promising residue-free alternative to traditional additives.

## 1. Introduction

To meet growing market demand for aquaculture products and to maximize production, feed additives such as growth promoters, vitamins, and antibiotics are commonly used in aquatic animal feed to combat pathogens and improve disease resistance (Banerjee and Ray, 2017). Growing concerns over antibiotic residues in aquaculture products have driven demand for additiveLfree alternatives. Therefore, aquaculture continues to explore new strategies for improved fish health and enhanced productivity. Probiotics are living microorganisms that, when administered in adequate amounts, confer beneficial effects to the host (Reid et al., 2019). Their use in aquaculture has shown promise as a very large number of genes and their products provided by microbial communities residing in water, sediment, and hosts (fish) are likely to impact aquaculture production, as summarized by Banerjee and Ray (2017), El-Saadony et al. (2021), and Rahayu et al. (2024). The direct impacts include: (i) improving the general health of the fish, enhancing the fish’s own digestive enzymes, (ii) enhancing feed efficiency, thus resulting in overall better growth performance, (iii) increasing resistance against infections, and (iv) improving reproduction. Aquaculture production is also likely to be indirectly impacted by improving water quality, reducing the abundance of pathogenic microbial species, providing nutrients for fish, and breaking down organic waste. In this context, probiotics offer a safe alternative to antibiotics and supplementing probiotics into water and feed has emerged as a strategy to influence microbial composition.

The common carp (*Cyprinus carpio*) is an important aquaculture species in many countries, especially in Eastern and Central Europe. In 2020, it was one of the most farmed fish species globally and (*The State of World Fisheries and Aquaculture 2020*, 2020) the increased demand is predicted in the future. Although, as reviewed by Dawood and Koshio (2016), research on the impact of probiotic supplementation in common carp has been intensively carried out for almost two decades, there has been little research related to the practical, on-farm (i.e., not laboratory) application of probiotic products, especially for probiotics that are mixed microorganism communities rather than specific bacterial species. The effect of feed supplementation with particular bacterial species on carp immune response and disease resistance has previously been studied and included *Lactobacillus casei* (Safari et al., 2022), *Lactobacillus fermentum* (Ahmadifar et al., 2019), and *Pediococcus acidilactici* (Hoseinifar et al., 2019). Furthermore, the effect of *Lactobacillus acidophilus* has been studied for multiple traits scored for common carp, quantifying growth performance and the level of immune response (Adeshina et al., 2020). Feng et al. (2022) and Gabr et al. (2023) reported a positive effect of *Lactobacillus acidophilus* on the growth rate and immune response of the carp. The improved immune response due to supplementation with *Enterobacter asburiae* was demonstrated by Li et al. (2023). Beneficial effects on the digestive system after *Lactobacillus rhamnosus* supplementation were reported by Chen et al. (2023), while Xia et al. (2024) observed an overall increased resistance to disease after supplementation of carp diet with *Bacillus subtilis*. Jwher and Al-Sarhan (2022) studied the general effect of probiotic supplementation on carp growth performance. However, in all the studies mentioned above, water tanks were used as laboratory conditions, and, to our knowledge, the only reported field experiment involving common carp probiotic supplementation was described by Mohammadian et al. (2022). It is important to realise that although laboratory-based studies in tanks provide a valuable and cost-effective initial experimental setup, they miss the considerable proportion of biological and environmental factors that are present in the field. Therefore, experiments conducted in real-world, farm settings provide more representative results and have greater implications for practical applications in aquaculture.

Our study was carried out as a field experiment in earthen ponds to reflect practical fish rearing conditions while investigating how supplementation with effective microorganisms affects microbial diversity, composition of the microbiome, and its dynamics over time. Our objectives were to assess the effects of supplementation across three different environments: (i) pond water, (ii) pond sediment, and (iii) fish gut, as well as its impact on fish growth performance. This was further enhanced by studying the effects of microbial supplementation on the growth performance of fish. We supplemented common carp (*Cyprinus carpio*) feed and ponds with effective microorganisms in a 103-day experiment and evaluated microbial composition. The microbiome composition was identified by sequencing the V3 and V4 hypervariable regions of the gene encoding the 16S rRNA ribosomal subunit and the metatranscriptome of water assessed by whole genome shotgun reads.

## 2. Material and methods

### 2.1. Experimental design

The experimental aquaculture facility consisted of 25 earthen ponds in the Gołysz experimental facility of the Polish Academy of Sciences, each covering an area of 600 m^2^ and supplied with freshwater from the Vistula River to maintain consistent physicochemical properties. The experiment spanned the entire fish production season, from May to October 2022, including a preliminary period of pond preparation setup from March to May. Experimental ponds were stocked by common carp fry at weight of 1 g at a density of 10,000 fry per hectare. The experimental ponds were classified into five groups, each consisting of five ponds. **Group 1** (control) did not receive supplementation. **Group 2** represented ponds with water supplementation with the supplement W1, without feed supplementation. **Group 3** represented ponds with water supplementation with the supplement W1 and feed supplementation with the supplement F1. **Group 4** represented ponds with water supplementation with the supplement W2, without any feed supplementation. **Group 5** represented ponds with water supplementation with the supplement W2 and feed supplementation with the supplement F2. According to the manufacturer’s information, the W1 water supplement contained *Lactobacillus sp*., three species of *Bifidobacterium sp*., *Carnobacterium divergens*, *Streptococcus salivarius*, and *Saccharomyces cerevisiae*. The feed supplement F1 contained all the above species and *Lactobacillus delbrueckii*, *Bacillus subtilis*, *Rhodopseudomonas palustris*, and *Rhodobacter sphaeroides*. The water supplement W2 and the feed supplement F2 consisted of *Lactobacillus plantarum, Lactobacillus casei*, and *Saccharomyces cerevisiae* along with several species that were confidentially protected by the manufacturer. The water supplements were applied on bottom (20 l/ha) and on water surface (20 l/ha) four times during the first month of experiment before first water and sediment sampling. The feed supplementation was mixed with the commercial grower feed Aller Classic Vitamax (Aller Aqua, Poland) and applied twice a week throughout the experiment at a concentration of 2 l/t. The detailed experimental design, including the supplementation schemes and available data types (described below), is presented in Fig. **1**.

**Fig. 1.**
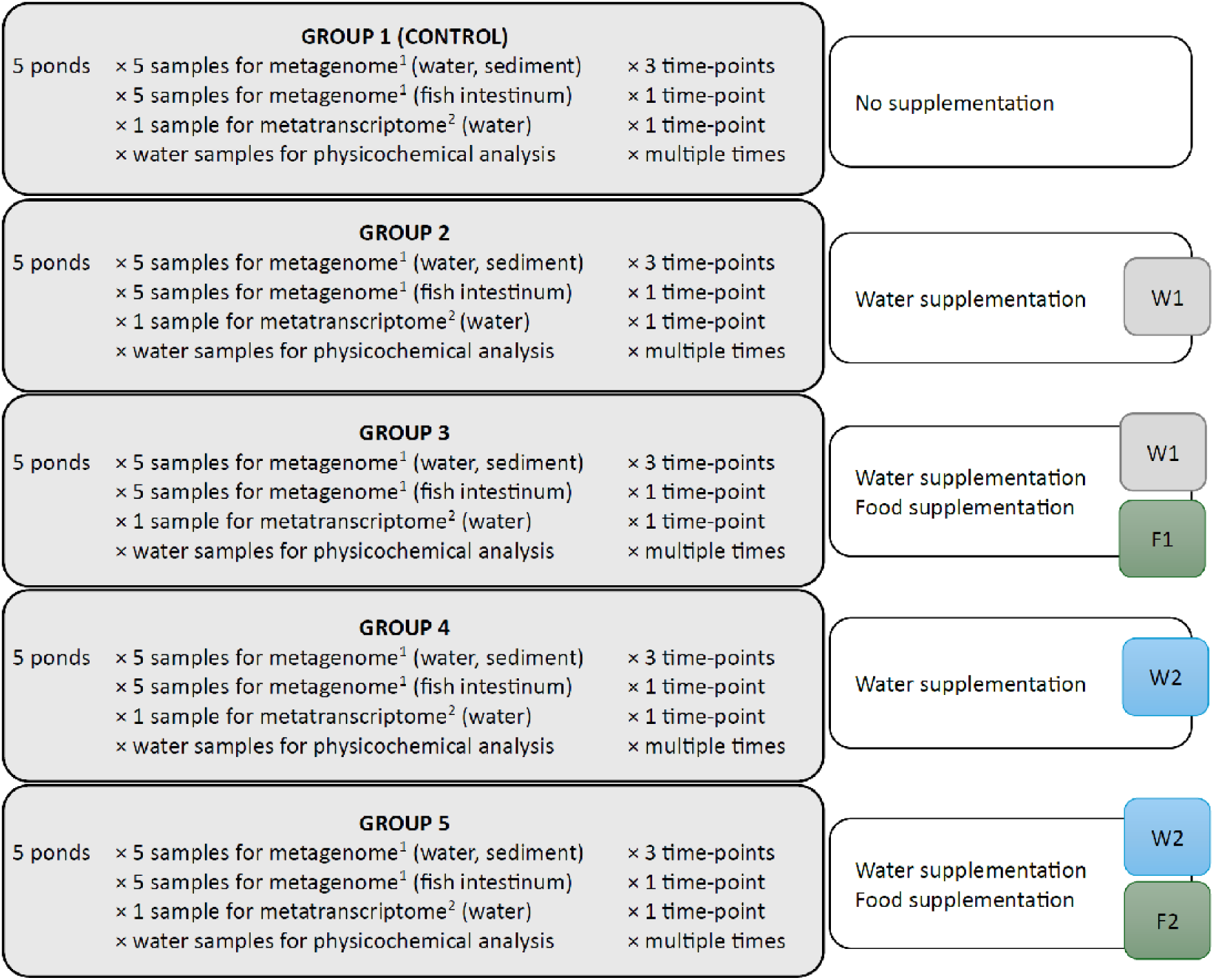
The experimental design includes 5 ponds without (Group 1), water (Groups 2 and 4) or water and feed (Groups 3 and 5) supplementation. Water supplements were represented by W1 and W2, whereas feed supplements were represented by F1 and F2. (1) represents 16S rRNA gene sequencing and (2) represents whole-sample RNA sequencing (metatranscriptome).

### 2.2. Sample preparation

#### 2.2.1. Sample collection

From each pond, three samples of water were collected three times during the experiment, specifically at the beginning 5 days after supplementation (T1), at the mid time-point 42 days after the 1^st^ sampling (T2), and at the end month of the experiment 54 days after the 2^nd^ sampling (T3). Sediment samples were collected twice, at the beginning and during the last month of the experiment, that is, 96 days after the 1^st^ sampling. Moreover, between days 101 and 103 after the beginning of the experiment, intestinal samples were collected from five common carp individuals from each pond (Fig. **1**).

The water samples were collected from the pond shoreline at an approximate depth of 0.5 m, each sampling site using a 2 m long and 1000 ml volume sampling pole. Three sampling sites were selected from the middle of the three shorelines of each pond. From each sampling site, equal volumes of water were collected, mixed, and stored in a 500 ml PEHD sterile bottle on ice and transported to the laboratory where they were stored at -20 °C until filtration. Microorganisms were collected by filtering approximately 500 ml of water through a sterile mixed cellulose esters (MCE) 0.22 μm pore size membrane (47 mm diameter, ChemLand, Poland). All filter membranes were stored at -80 °C until DNA extraction. Sediment samples were collected using the three-point sampling method. Sediments were sampled using a corer made of 5.5 cm diameter metal tubing attached to a 2 m. long wooden pole. The samples were collected from the surface of the collecting corer using a sterile metal spoon and a 50 ml sterile falcon tube. Approximately 10 g of sediment was collected from three different sampling sites (the same as water was collected) in each pond and mixed immediately after collection to form a single sample.

#### 2.2.2. 16S rRNA sequencing

Commercial DNA isolation kits, dedicated to environmental samples, were used for DNA isolation. The PowerSoil®, the DNeasy PowerWater Kit as well as the QIAamp PowerFecal Pro DNA Kit (QIAGEN®, Germany), were applied for sediment, water, and intestine microbiota respectively. The quality and concentration of DNA was assessed by gel electrophoresis and fluorometry. DNA samples were normalized to a concentration of five ng/μl. Then, 20 cycles of PCR were performed using primers containing tag sequences. Electrophoresis of the PCR products was performed on a 1.5% agarose gel (Invitrogen, cat. no. 16500500) stained with GelGreen dye (Biotium cat. no. #41004). The samples were loaded with Orange G loading dye (6X) (Thermo Scientific, cat.no.J60562.AC). The remaining 9 ul of samples were supplemented with MQ water to 20 ul and purified using the SPRI method, using KAPA Pure Beads reagent (KAPA Biosystems cat. no. 07983271001) in the ratio sample: reagent 1:0.8. Libraries were constructed from the prepared amplicons by incorporation of adapters IDT for Illumina Nextera DNA UD Indexes Set A (96 Indexes, 96 samples) (Illumina, cat.no.20027213) in 8 PCR cycles. KAPA HiFi HS RM was used (KAPA Biosystems cat.no.795893500) according to the manufacturer’s protocol. The libraries obtained were collected in 11 pools of 13-24 libraries each. The pools were surrendered for fragment length control using the Tape Station 2200 analyzer and a set of High Sensitivity D5000 Reagents (Agilent, cat.no.5067-5593). The concentration of the libraries was determined by qPCR using the Kapa Library Quantification kit, according to the manufacturer’s instructions (Kapa Biosciences, cat.no.KK4824). Fragment selection was carried out using Kapa Pure Beads in one-step purification. Sequencing of selected hypervariable regions (V3 and V4) of the gene encoding the 16S subunit of a ribosome (16S rRNA gene) was performed on an Illumina NovaSeq 6000 instrument using the NovaSeq 6000 SP Reagent Kit v 1.5 (500 cycles), with pair-end reads of 2×250 cycles according to the manufacturer’s instructions with 1% addition of the Phix control library (Illumina, code no. FC-110-3001).

#### 2.2.3. Metatranscriptome sequencing

One water sample per pond was collected for metatranscriptome analysis on 102-104^th^ day of–experiment. For RNA isolation, water samples were filtered using a sterile vacuum system through sterile mixed cellulose esters (MCE) 0.22 μm pore size membranes (47 mm diameter, ChemLand, Poland). The membranes were placed in sterile Eppendorf tube and immediately freezed in liquid nitrogen freeze to preserve the degradation of the sample RNA and stored at -80 °C. The RNeasy PowerWater Kit (QIAGEN®, Germany) was used according to the manufacturer’s instructions. RNA samples were qualitatively analyzed using an Agilent TapeStation 2200 analyzer and High Sensitivity RNA ScreenTape reagents (Agilent, no.5067-5579), followed by eukaryotic ribodepletion using the NEBNext® Globin & rRNA kit Depletion Kit (Human/Mouse/Rat) (New England Biolabs, cat. no. E7750X) and library construction using the KAPA RNA HyperPrep Kit (KAPA Biosystems, cat.no.8098107702), according to the manufacturer’s recommendations, starting with 255 ng of material. The KAPA_UDI_Indexes adapters were used (96 indices, 96 samples) (KAPA Biosystems, cat. no. 08098140702). RNA was fragmented at 85 °C for 6 minutes and ten cycles of enrichment by amplification were performed. The libraries obtained were subjected to fragment length control using the Agilent TapeStation 2200 analyzer and the High Sensitivity D1000 Reagents set (Agilent, cat.no.5067-5585). The concentration of libraries was determined by qPCR using the Kapa Library Quantification kit (Kapa Biosciences, cat.no.KK4824). The manufacturer’s protocols were used. Sequencing was performed on an Illumina NovaSeq 6000 instrument using NovaSeq 6000 SP Reagent Kit v 1.5 (500 cycles), with pair-end reads of 2×250 cycles according to the manufacturer’s instructions with 1% addition of the Phix control library (Illumina, cat.no.FC-110-3001). Due to poor data quality, the two control samples were discarded.

### 2.3. Phenotypic data

Phenotypic data were collected from 750 individuals (30 fish per pond) on the last day of the experiment. Weight, total length (including tailfin), and pre-dorsal height were measured. Weight was measured with an accuracy of 0.1g. Length and height measurements were made on graph paper with an accuracy of 0.1mm.

### 2.4. Bioinformatic analysis

#### 2.4.1. 16S rRNA

The processing and analysis of 16S rRNA sequences were performed using QIIME2 software (Bolyen et al., 2019). The processing pipeline involved: (1) the assessment of raw read quality, (2) trimming sequences from 5’ and 3’ ends with quality scores below 30, (3) removing sequencing adapters, (4) removing sequences shorter than 200 bp, (5) merging paired end reads, (6) additional quality filtering step (quality filter q-score) and (7) denoising (qiime deblur denoise-16S). This pipeline yielded Amplicon Sequence Variants (ASVs) that were used to determine the microbial composition of the samples. The taxonomic assignment was completed using QIMME2 feature classifier implementing a Bayesian classification model pre-trained on 16S rRNA data. All analyses following taxonomic assignment were performed using the SILVA (v.138.1) database (Quast et al., 2012) at the family taxonomic level.

The alpha diversity metric quantifying microbial diversity within a pond was quantified using Shannon’s index: 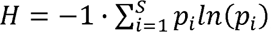, where *S* represents the number of families present in a given pond and *p_i_* is the frequency of the i-th family. Bacterial composition of water, sediment, and gut environments was expressed as PCA-based embedding and then visualised using the first two principal component vectors. For the computation of embeddings, the *prcomp* function from the *stats* library implemented in R (2016) was applied with default parameters, using abundances normalised with Centered Log-Ratio (CLR) transformation: 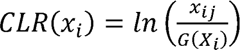, where *x_ij_* represents the raw abundance of *j*-th bacterial family within *i*-th sample (fish or pond) and *G*(*X_i_*) is the geometric mean of raw abundances representing sample *i* (Aitchison, 1982).

Furthermore, we analysed the differences in abundance of particular bacterial families, using the DESeq2 package (Love et al., 2014) implemented in R. For gut samples, pairwise comparisons were made between the control (**Group 1**) and each experimental group (**Group 2**, **Group 3, Group 4**, or **Group 5**). For environmental samples, comparisons were made at three (water) and two (sediment) time points. At the first time point, the control (**Group 1**) was compared to all experimental groups combined (**Group 2** - **Group 5**). At the second and third time points, as for gut samples, pairwise comparisons were made between the control group (**Group 1**) and each individual experimental group (**Group 2**, **Group 3**, **Group 4**, or **Group 5**). Bacterial family abundances were normalised using the default median-of-ratios normalisation implemented in DESeq2 (Anders and Huber, 2010). Then the differential abundance was expressed as the log_2_FoldChange (LFC) between samples, while hypotheses testing was based on the Wald test 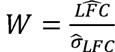, which under the null hypothesis of no differences in abundances asymptotically follows the standard normal distribution. The nominal P-values corresponding to each tested family were corrected for multiple testing using the Bonferroni approach.

#### 2.4.2. Metatranscriptome

Metatranscriptomes were primarily used to derive unbiased taxonomic compositions of water samples and were processed using the MG-TK pipeline (Anthony) run in a reference-based mode. Basic filtering, functional and phylogenetic profiling are an adapted protocol of the methods used in , 2021). Filtering based on read quality was performed with a simple demultiplexer (Hildebrand et al., 2014) by removing: (i) reads with estimated accumulated error exceeding 2.5 with probability ≥ 0.01 (Trivedi et al., 2016) (ii) reads with more than one ambiguous position, or (iii) reads containing a homonucleotide run longer thanL15Lbp. Reads were trimmed if the base quality dropped below 20 in a window of 18 bases at the 3′ end, or if the accumulated error exceeded 1. The filtered reads were profiled using the miTag approach (Salazar et al., 2021). The reads corresponding to the rRNA subunits were extracted using SortMeRNA (Kopylova et al., 2012) and then searched against the SILVA (v.128) database (Quast et al., 2012). Reads matching the database with ELvalues <L10^−4^ were further filtered with custom Perl and C++ scripts using FLASH (Magoč and Salzberg, 2011) to merge all matched read pairs. In case read pairs could not be merged, single reads were interleaved so that the second read pair was reverse complemented and then sequentially added to the first read. The candidate interleaved or merged reads were then fine-matched to the SILVA database using Lambda3 (Hauswedell et al., 2024). To determine the identity of filtered reads based on Lambda3 matches, the lowest common ancestor (LCA) algorithm adapted from LotuS (Hildebrand et al., 2014) was applied. The abundance of each taxon was then normalized by dividing it by the total number of reads per sample to account for uneven sequencing depth across samples.

### 2.5. Modelling fish growth

The association between microbial family abundance in the gut and the phenotypic measurements of the individual fish (weight, height, and length) was analysed using two-way hierarchical analysis of variance (ANOVA) and a linear penalized regression. In the ANOVA, an experimental group was used as the class variable, and ponds were modelled as nested effects. For linear regression, the data preprocessing step comprised averaging the CLR normalised abundance of each bacterial family within each pond over the five fish for which 16S rRNA sequences were available. Then, the LASSO linear regression (Tibshirani, 1996) that employs L1 norm regularization was fitted by minimizing:

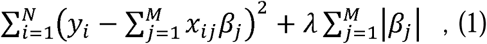

where *y_i_* is the phenotypic measurement for *i*-th fish, *x_ij_* represents the abundance of *j*-th bacterial family averaged over the five ponds from experimental conditions corresponding to fish *i*, /*β_j_* is the effect of *j*-th family, *λ* is the regularisation parameter, *N* is the number of individuals with phenotypes, and *M* is the overall number of bacterial families. Note that the hyperparameter *λ* was arbitrarily set to 0.1 by comparing different *λ* estimates from a cross validation approach implemented in the *glmnet* R package.

## 3. Results

### 3.1. Within sample diversity

The median microbial diversity of the water samples (Fig. **2**) at the first sampling time-point (T1) was similar. However, the ponds supplemented with water supplement W1 were more uniform in their alpha diversity (standard deviations 0.81 and 0.80) than the five control ponds (standard deviation 2.32) and the ponds supplemented with water supplement W2 (standard deviations 2.52 and 1.61). The dynamics of within-pond diversity exhibited a distinct pattern that differed between the control and the experimental ponds. In the control ponds, the microbial diversity increased with time, while all experimental points demonstrated an initial increase in diversity from T1 to T2, followed by a decrease towards the end of the experiment (T3). Sediments generally exhibited less pronounced changes in alpha diversity over time compared to water samples. Although no strong patterns emerged in the diversity attributed to different groups, in general, the alpha diversity of the experimental groups tended to increase with time, while the opposite was observed for the control group (Fig. **2**). When examining individual ponds serving as experimental replicates (as shown in Fig. **3**), changes in sediment diversity over time appeared more pronounced in the control ponds than in the supplemented ponds. In particular, in four out of five control ponds, the diversity decreased within the range of 1.48 to 5.72, whereas in a single pond, an opposite trend was observed. The diversity dynamics of supplemented ponds revealed no pattern regarding increase or decrease in diversity within each pond, varying in a range 0.20-2.78 (for decreased diversity) and 0.22-3.24 (for increased diversity). Regarding the microbial communities in the fish gut visualized in Fig. **4**, our analysis revealed that its diversity was markedly lower than that of the environmental samples (water and sediment). Fish representing **Group 3** receiving water and feed supplementation W1-F1 exhibited slightly higher diversity levels (average of 4.65) compared to 3.84, 3.03, 3.82, and 3.83 in the other groups. However, as expressed by within-group standard deviation of individual alpha diversities, the level of gut microbiome diversity was more similar among individuals subjected only to water supplementation in **Group 2** (1.08) and water and feed supplementation W2-F2 supplementation in **Group 5** (1.04). The alpha diversity of sequenced supplements was considerably lower than that of gut and environmental diversity, amounting to 0.57, 0.63, and 0.38, respectively for W1-F1, and commonly for W2-F2, which constitutes the same product supplemented in two different forms to water and feed. More specifically, W1 comprised two families with relative abundance of: *Lactobacillaceae* (0.976) and *Acetobacteraceae* (0.024), F1 contained three families of the relative abundance of: *Lactobacillaceae* (0.900), *Clostridiaceae* (0.093), and *Sporolactobacillaceae* (0.007), W2-F2 consisted of *Clostridiaceae* (0.909), *Lactobacillaceae* (0.055), and *Paenibacillaceae* (0.036).

**Fig. 2.**
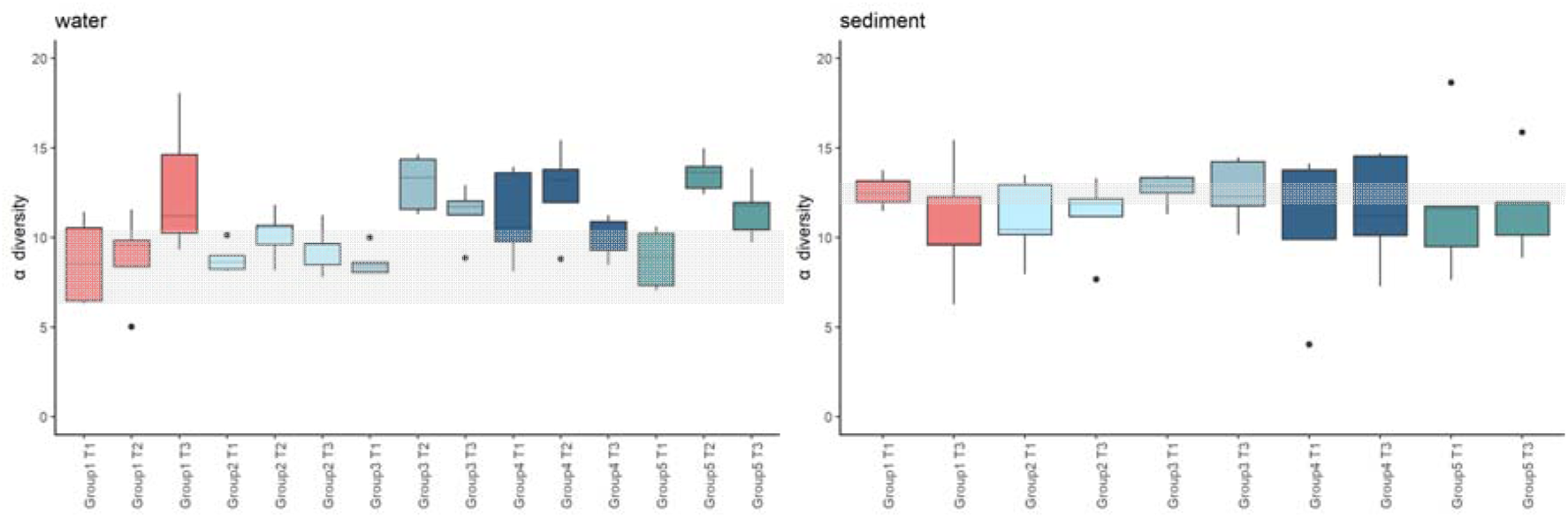
Alpha diversity (Shannon index) of water and sediment samples. T1 (day 5), T2 (day 47), and T3 (day 100) represent consecutive time-points of sampling. Group 1 is the control group (5 ponds), Group 2 is experimental group W1 (5 ponds), Group 3 is experimental group W1-F1 (5 ponds), Group 4 is experimental group W2 (5 ponds), and Group 5 is experimental group W2-F2 (5 ponds), where F and W denote feed and water supplement respectively. The shaded areas mark 75% of the variation in alpha diversity observed in the control ponds at T1.

**Fig. 3.**
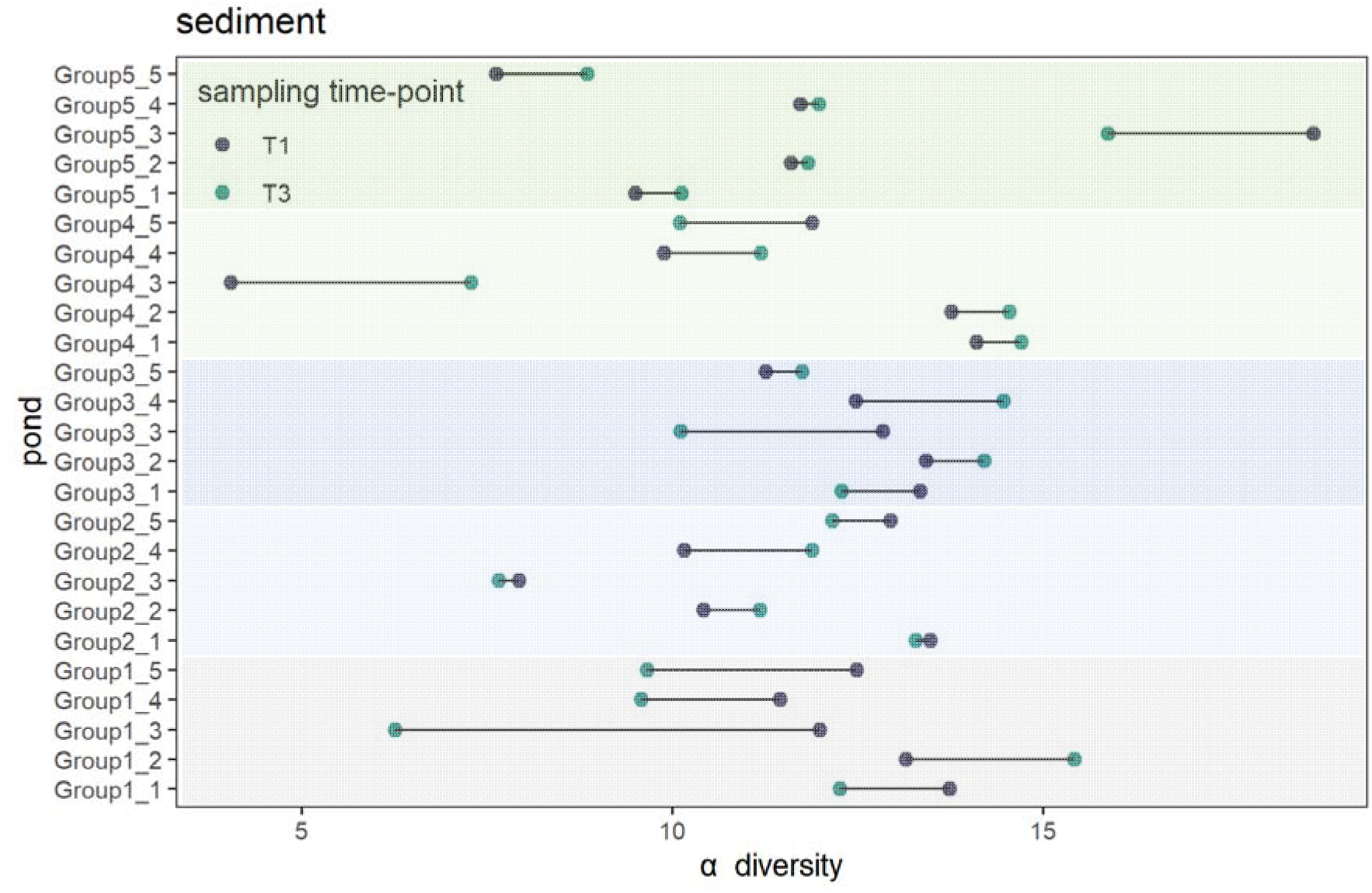
Alpha diversity (Shannon index) of sediment samples. T1 (day 5) and T3 (day 100) represent consecutive time-points of sampling. Group 1 is the control group (5 ponds), Group 2 is experimental group W1 (5 ponds), Group 3 is experimental group W1-F1 (5 ponds), Group 4 is experimental group W2 (5 ponds), and Group 5 is experimental group W2-F2 (5 ponds), where F and W denote feed and water supplement respectively.

**Fig. 4.**
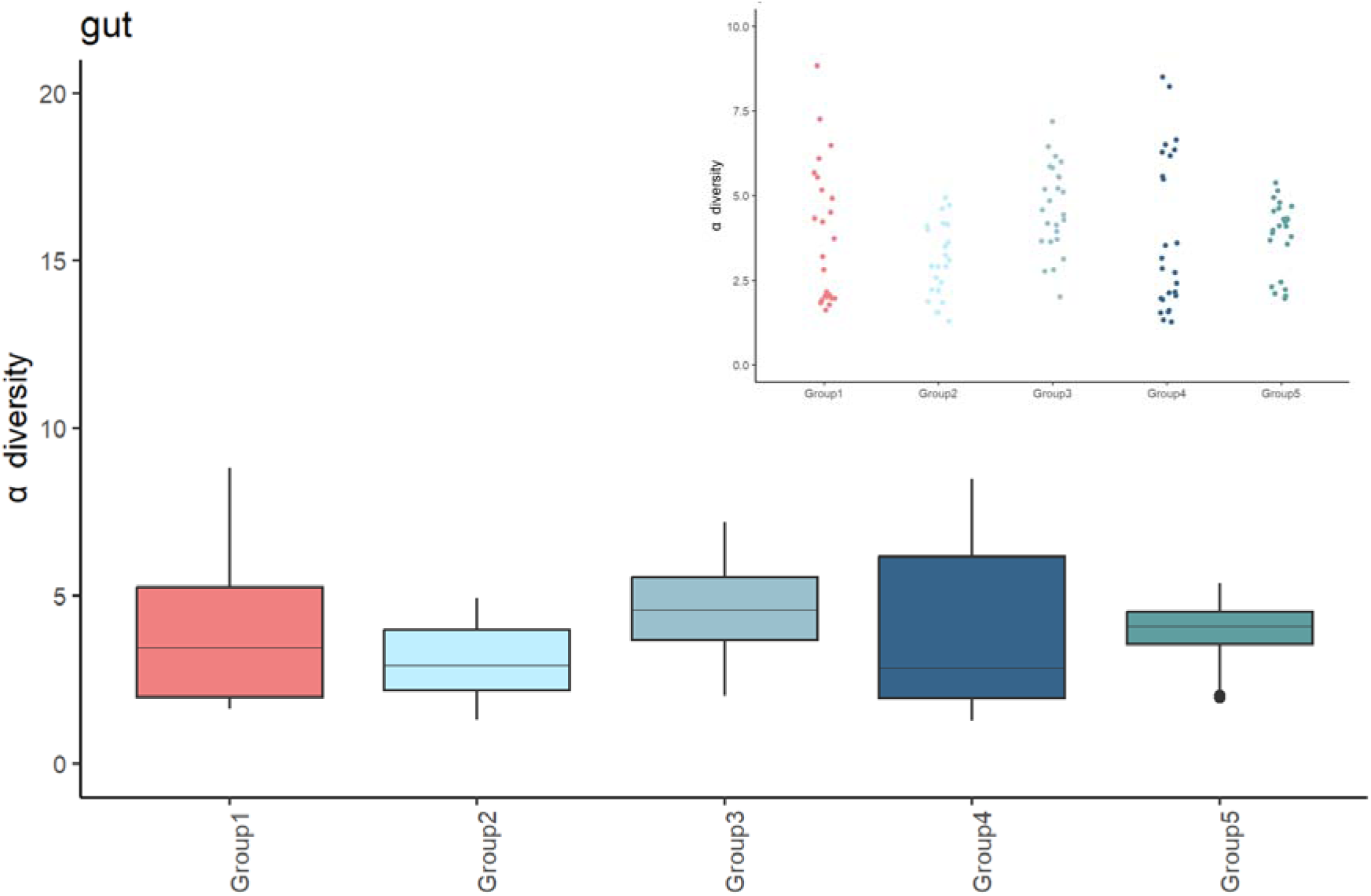
Alpha diversity (Shannon index) of gut samples for ponds and the individual fish (inset). Group 1 is the control group (5 ponds), Group 2 is experimental group W1 (5 ponds), Group 3 is experimental group W1-F1 (5 ponds), Group 4 is experimental group W2 (5 ponds), and Group 5 is experimental group W2-F2 (5 ponds), where F and W denote feed and water supplement respectively.

### 3.2. Dominant bacteria families

In water, *Comamonadaceae* was consistently the most abundant family under all experimental conditions. *Burkholderiaceae*, *Chitinophagaceae*, and *Methylophilaceae* were also highly abundant in most ponds. Notably, bacterial families provided in the supplements were either undetectable or of very low abundance, with the exception of *Paenibacillaceae* from the W2 and F2 supplements, which was highly abundant in one pond from experimental Group 5 supplemented by W2-F2, but somewhat unexpectedly in one pond from experimental Group 3, which was not supplemented by W2 or F2 (Fig. **5**).

**Fig. 5.**
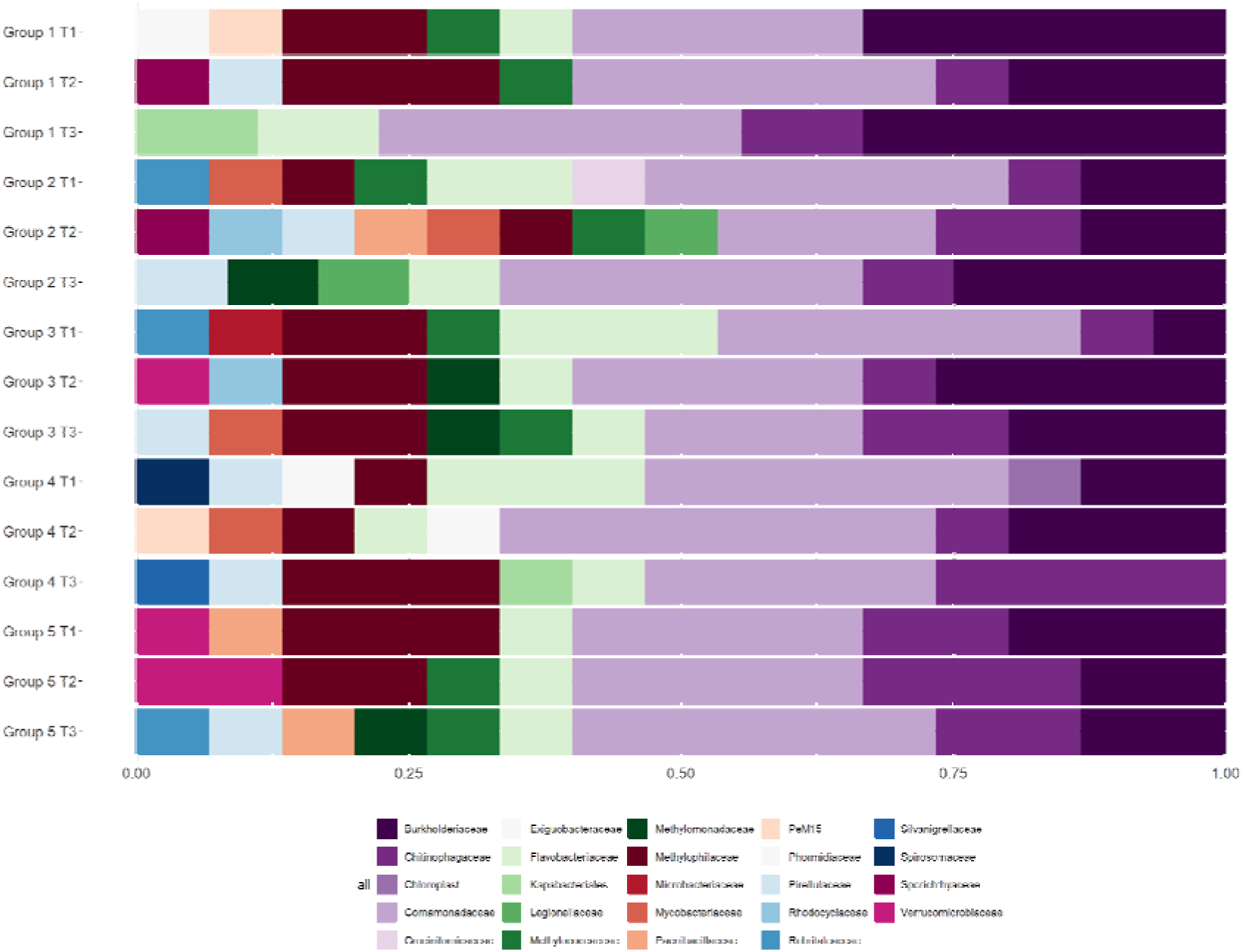
Frequency of the three most abundant families in water pooled over the experimental groups. T1 (day 5), T2 (day 47), and T3 (day 100) represent consecutive time-points of sampling. Group 1 is the control group (5 ponds), Group 2 is experimental group W1 (5 ponds), Group 3 is experimental group W1-F1 (5 ponds), Group 4 is experimental group W2 (5 ponds), and Group 5 is experimental group W2-F2 (5 ponds), where F and W denote feed and water supplement respectively.

Sediment samples demonstrated considerably higher variation in the most abundant families across ponds than water samples. They showed a variable pattern of high abundance, where numerous families were highly represented but often uniquely within a single sample (Fig. **6**). This may have been caused by the greater influence of spatial sampling heterogeneity within the ponds, where the water exhibited greater spatial uniformity owing to increased movement within a pond. *Bacteroidetes_vadinHA17* was present in most, albeit not all, ponds, while each group (including the control) contained relatively high counts of *Bacteroidetes_vadinHA17*, *Rhodocyclaceae*, *Nocardioidaceae*, and *Phormidiaceae*. Supplementation with W2/W2-F2 resulted in more variable patterns of highly abundant bacterial families, whereas the most consistent pattern was observed in five ponds supplemented with W1. Most of the bacterial families introduced with supplements were not established in the sediment, with the exception of *Clostridiaceae*, which was present in almost all ponds, including the control (i.e., unsupplemented ponds), indicating that it is a naturally abundant bacterial family. The abundance of *Paenibacillaceae* supplemented with W2-F2 was detected in only one control pond and one pond supplemented with W2.

**Fig. 6.**
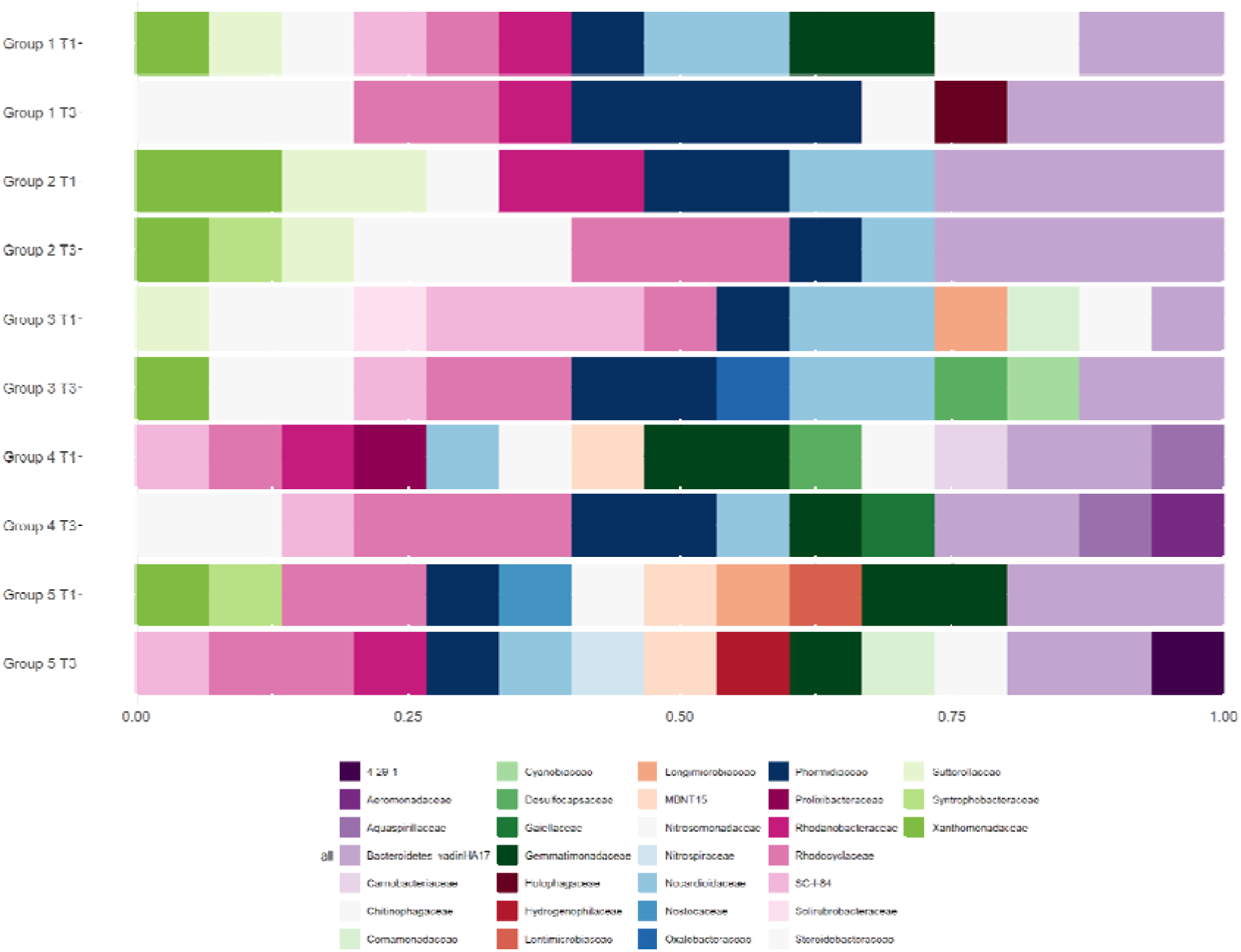
Frequency of the three most abundant families in sediment pooled over the experimental groups. T1 (day 5) and T3 (day 100) represent consecutive time-points of sampling. Group 1 is the control group (5 ponds), Group 2 is experimental group W1 (5 ponds), Group 3 is experimental group W1-F1 (5 ponds), Group 4 is experimental group W2 (5 ponds), and Group 5 is experimental group W2-F2 (5 ponds), where F and W denote feed and water supplement respectively.

In the gut, *Aeromonadaceae* was among the most abundant families in each fish, except for a single individual. Furthermore, *Erysipelotrichaceae* and *Fusobacteriaceae* were among the most common families regardless of the experimental conditions (Fig. **7**). Also , the within-pond similarity among fish regarding the most abundant families was not higher than among fish from different ponds. Regarding the bacterial families contained in the supplements, *Clostridiaceae* that was supplemented in F1 and F2 was detected in most individuals, except nine fish, but the pattern was not related to supplementation, indicating that the family is naturally abundant in the gut, with no supplementation required. Unexpectedly, *Paenibacillaceae* supplemented with F2 was present in two individuals, but none of them were supplemented with F2. The other families supplemented in feed (*Lactobacillaceae* and *Sporolactobacillaceae*) were not present. Evidence indicates that feed supplementation has little to no effect on gut microbiome composition.

**Fig. 7.**
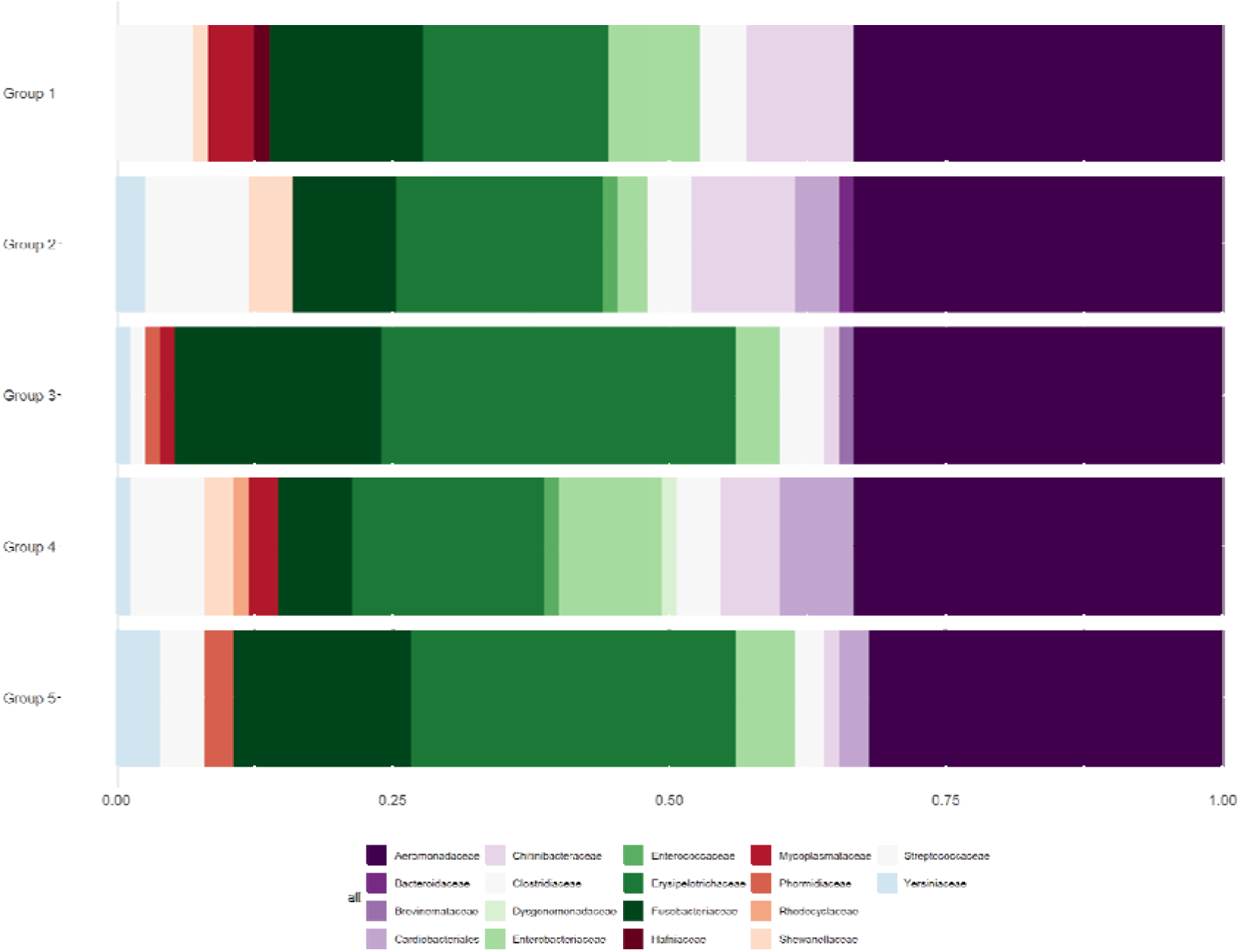
The frequency of the three most abundant families in the fish gut pooled over the experimental groups. Group 1 is the control group (5 ponds), Group 2 is experimental group W1 (5 ponds), Group 3 is experimental group W1-F1 (5 ponds), Group 4 is experimental group W2 (5 ponds), and Group 5 is experimental group W2-F2 (5 ponds), where F and W denote feed and water supplement respectively.

### 3.3. Differences in overall bacterial composition

Fig. **8** summarises the visual representation of the bacterial family abundances characteristic to each of the studied environments (water, sediment, and gut). Although no distinct clustering patterns related to the experimental conditions emerged, it is clear that within the course of the experiment, both water and sediment bacterial communities tended to converge, becoming more similar across experimental conditions.

**Fig. 8.**
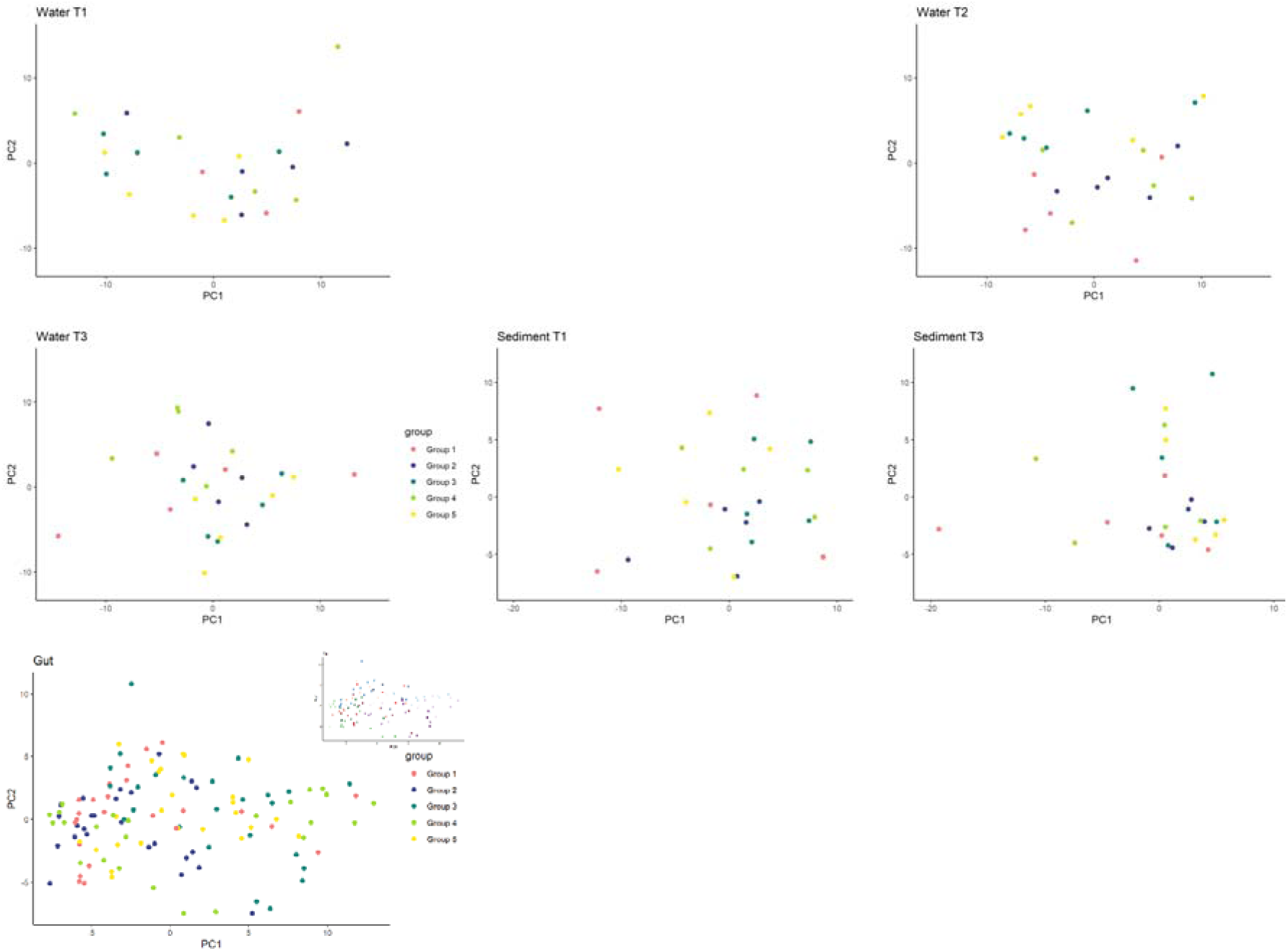
PCA representation of bacterial family abundance characteristic for each pond. T1 (day 5), T2 (day 47), and T3 (day 100) represent consecutive time-points of sampling. Group 1 is the control group (5 ponds), Group 2 is experimental group W1 (5 ponds), Group 3 is experimental group W1-F1 (5 ponds), Group 4 is experimental group W2 (5 ponds), and Group 5 is experimental group W2-F2 (5 ponds), where F and W denote feed and water supplement respectively.

### 3.4. Differences in abundance of single families

Since no clear pattern emerged for overall microbial diversity, the second part of the study focused on particular families.

The analysis of differential abundances of bacterial families in water samples at the first sampling time-point revealed no significant differences in abundances between the control ponds and the experimental ponds, even though T1 sample collection was one week after the first water supplementation. At the 2nd sampling time-point, there were still no significantly altered abundances between the control group (**Group 1**) and **Group 2**, but in the remaining three supplemented groups a significant increase in the abundance of *Rubritaleaceae* was observed. The corresponding LFCs were 6.4 (P=0.003), 7.5 (P=1·10^-4^), and 7.7 (P=7·10^-5^), respectively for **Groups 3-5**. Furthermore, a significant increase in *Solirubrobacteraceae* abundance with LFC of 7.6 (P=0.022) was detected in **Group 5**. However, the observed effect did not persist toward the end of the experiment, hence in the last sampling time-point (T3) which occurred 54 days after T2, the differences in abundance of *Rubritaleaceae* and *Solirubrobacteraceae* were no longer significant. Still, the microbial community demonstrated further dynamics as at the last time-point, significant decreases in the abundances of bacterial families were observed involving *Carnobacteriaceae* in **Groups 2** and **3** with LFCs -21.6 (P=2·10^-12^) and -22.0 (P=1·10^-13^) respectively, *Geobacteraceae* (LFC=-22.7, P=7·10^-14^) in **Group 2**, *Sphingobacteriaceae* (LFC=-7.6, P=0.047) in **Group 3**, *Methylomonadaceae* (LFC=-8.3, P=0.007) and *Moraxellaceae* (LFC=-22.7, P=8·10^-14^) in **Group 4**, as well as *Exiguobacteraceae* (LFC=-8.8, P=0.037) in **Group 5**. Note that, except *for Methylomonadaceae*, none of these families was among the most abundant, indicating that the supplementation predominantly acted on the less predominant families.

The above results obtained for T3 were further refined with the metatranscriptome data, allowing for an unbiased view of both the eukaryotic and bacterial microbiomes, which were inaccessible through our 16S rRNA gene amplicon sequencing. In total, 231,353,334 metatranscriptome reads from 533,108,406 were filtered out during quality control. On average, 2,138,017 and 1,168,061 reads per pond matched the large ribosomal subunit (LSU) and the small ribosomal subunit (SSU), respectively, and were taxonomically assigned. Regarding LSU-based annotations, no significant differential abundance was detected between the control and experimental ponds, which was presumably due to the poorer composition of the SILVA database for LSU sequences. Annotation based on SSU indicated a significant differential abundance of non-bacterial microorganisms. Two eukaryotic genera: a monotypic fungi *Linostomella* sp in **Group 3** (LFC=5.3, P=2·10^-4^) and **Group 4** (LFC=9.8, P=9·10^-4^) as well as an unicellular protist *Dileptus* sp. in **Group 2** (LFC=22.2, P=1·10^-13^) were significantly more abundant in experimental ponds, while the abundance of another protist species, *Paramecium bursaria*, decreased significantly under W2 supplementation in **Group 4** (LFC=-7.3, P=0.045).

For sediment samples, at the first time-point, a significant increase in abundance was detected after supplementation for **Group 3**, with *Cyanobiaceae* (LFC = 8.4, P = 0.012), which even became one of the three most abundant families in a pond representing this experimental group. At the second time-point, bacteria representing the *Flavobacteriaceae* family (LFC=-7.1, P=0.039) significantly decreased abundance in experimental ponds representing **Group 3**, while the abundance of *Streptococcaceae* (LFC=22.9, P=1·10^-16^) and *Aquaspirillaceae* (LFC=25.0, P=6·10^-23^) significantly increased in **Group 4**. The latter was one of the most abundant families in the two ponds of this group.

In the fish gut, supplementation led to an increase in several bacterial families compared with the control ponds. Supplementation with W1-F1 (**Group 3**) resulted in a significant increase in the abundance of *Weeksellaceae* (LFC=6.5, P=2·10^-7^), *Geobacteraceae* (LFC=6.8, P=0.001), and *Methylomonadaceae* (LFC=8.9, P=4·10^-14^). Water supplementation with W2 in **Group 4** led to a significant increase in the abundance of *Erysipelotrichaceae* (LFC=2.0, P=0.008), which was already among the most abundant families in the gut microbiomes of many fish in all experimental ponds. Furthermore, in **Group 4**, we observed an increased abundance of *Yersiniaceae* (LFC=4.1, P=0.013), *Enterobacteriaceae* (LFC=6.5, P=1·10^-3^), and *Cardiobacteriales* (LFC=2.8, P=0.010) bacterial families that were common in the gut microbiomes of some fish from experimental ponds. Supplementation with W2-F2 in **Group 5** affected the abundance of the four families. A significant increase was detected for: *Erysipelotrichaceae* (LFC=2.4, P=2·10^-5^) which was a commonly abundant family in fish from all ponds, another commonly abundant family *Cardiobacteriales* (LFC=2.4, P=0.023), as well as for *Methylomonadaceae* (LFC=7.2, P=9·10^-7^) that was not among the most highly abundant. Conversely, *Streptococcaceae,* which was a highly abundant family in all ponds, significantly decreased in abundance (LFC=-3.6, P=0.032) after W2-F2 supplementation in **Group 5**.

Notably, **Group 2** supplemented with W1 showed no significant alterations in bacterial family abundance in either the sediment or gut environments.

### 3.5. Fish body measurements

For each of the three body measurements depicted in Fig. **9**, the differences between experimental groups were significant, varying between P=0.0105 for weight and P=7·10^-5^. For height. Specifically, fish in **Group 2** were significantly larger than those in the control group (on average: 4.39 g heavier, 0.42 cm longer, and 0.20 cm higher), but also more than in the experimental **Groups 3** and **4**. However, the within group variability between ponds was much more significant than the variability between groups.

**Fig. 9.** Fish body measurements. Group 1 is the control group (5 ponds), Group 2 is experimental group W1 (5 ponds), Group 3 is experimental group W1-F1 (5 ponds), Group 4 is experimental group W2 (5 ponds), and Group 5 is experimental group W2-F2 (5 ponds), where F and W denote feed and water supplement respectively.

The common feature of the penalized linear regression models was their poor fit expressed by R^2^ of 0.22 for weight, 0.10 for height, and 0.19 for length. When correlating microbial abundance and fish growth, we found that the highest R^2^ (0.22) and the largest number of bacterial families (23) were observed for weight. The highest positive effect of 8.3 g weight associated with an increase in one abundance unit was estimated for the order *Flavobacteriales* without a particular family assigned, and the highest negative effect of a 5.9 g decrease in weight was associated with an increased abundance of *Silvanigrellaceae*. For length, 13 bacterial families were selected with the highest positive effect of 0.6 cm was associated with the increased abundance of *Nocardiaceae* and the highest negative effect of 0.9 cm, with *Silvanigrellaceae*, i.e. the same family that also negatively influenced weight. *Nocardiaceae* also had a high positive association with weight. For height only six families were selected, albeit with overall low effects, ranging between 0.02 cm (*Sporomusaceae*) and -0.05 cm (*Phormidiaceae*).

## 4. Discussion

This study is novel in that it characterized three different environments, that is, water, sediment, and fish gut, to evaluate the efficacy of effective microorganisms supplementation in modifying the microbial composition and improving the growth of common carp in earthen ponds. The key observation from our study was that most of the supplemented bacterial families were not established in the studied environments. Moreover, the supplementation also did not markedly influence the overall composition of the studied environments. Most changes over time were observed for water, in contrast to the control ponds, where the natural diversity increased during the course of the experiment; in all supplemented groups, the diversity dynamics of the water microbiome were the same after the initial increase (that may be due to the supplementation), and there was a decrease in diversity towards the end of the experiment. This suggests that, although the introduced bacterial families may not have directly established themselves, they indirectly influenced the ecosystem by altering the abundance of other families, which, however, has only a temporary effect. Also Huerta-Rábago et al. (2019) observed an increase in water alpha diversity of untreated, control ponds and changes in alpha diversity in supplemented ponds, albeit following a different pattern than it was observed in our study. As demonstrated by PCA not only no distinct bacterial communities were established by supplementation, but also for water and sediment, during the course of the experiment the microbiomes tended to converge , i.e. be more similar across experimental conditions and the control. In addition, the abundance of families that were most highly represented in each environment was not markedly influenced by the supplementation.

However, some single families were differentially abundant. Species from the *Rubritaleaceae* family, whose abundance was increased in experimental **Groups 3-5** show antioxidative activities (Shindo et al., 2007) and are involved in polysaccharide degradation (Martinez-Garcia et al., 2012). The increased abundance of *Solirubrobacteraceae* was somewhat unexpected, since the family is rather typical for soil environments and may represent a sampling or sequencing artifact. *The abundance of Carnobacteriaceae* in water significantly decreased at the end of the experiment in **Groups 2** and **3** (both supplemented with F1), which represents the group of lactic acid bacteria that in aquaculture is predominantly found in fish distal intestine and exhibits a positive effect on digestion (Ringø et al., 2003). However, species representing this family have also been found in lakes. This family has also been investigated in the context of food degradation, preservation, and degradation of oils in wastewater processing (Rosenberg et al., 2014a). In addition, the abundance of *Geobacteraceae* was significantly altered by supplementation. In water, it decreased at the end of the experiment in **Group 2**, whereas in the gut, its abundance was higher after supplementation with F1. Species from this family predominantly reside in sediment, where they induce the release of soluble Fe(II) from insoluble Fe(III) into water, which may further stimulate the growth of other microorganisms (Rosenberg et al., 2014b). *Sphingobacteriaceae* that was less abundant in water of experimental **Group 3** originally represents soil inhabiting family, but is also reported in aquaculture as containing species that metabolite cellulose, chitin, collagen, and nitrogen compounds in water (Bartie et al., 2005). *Flavobacteriaceae,* which significantly decreased in abundance in sediment after W1-F1 supplementation in **Group 3**, are common in planktonic freshwater communities (Riemann and Winding, 2001) and are regarded as potentially pathogenic. Members of this family often metabolise large molecules, such as polysaccharides and proteins (Rosenberg et al., 2014c). *Weeksellaceae*, which was significantly increased in **Group 3**, represent an unwanted, pathogenic family that promotes inflammation and disrupts the formation of intestinal villi in fish (Liu et al., 2020). *Erysipelotrichaceae*, which is regarded as an unwanted, potentially pathogenic family, was not only among the most abundant in the fish gut, but also its gut abundance was significantly increased after F2 supplementation. The same was true for the significant increase in the pathogenic *Yersiniaceae* family. This observation contradicts recent results obtained for the shrimp *Penaeus vannamei* (Luo et al., 2024) and largemouth bass (Li et al., 2024). In contrast, **Group 4** supplemented with W2-F2 revealed a significant increase in the beneficial family of *Enterobacteriaceae,* which is considered a probiotic supplement (see, for example, Li et al. (2023)), and significantly decreased abundance of *Streptococcaceae* family that contains fish pathogenic species such as *Lactococcus piscium* (Rosenberg et al., 2014a).

### 4.1. Supplementation fish growth

Even though in our experiment no significant changes in water and sediment microbial communities due to the supplementation occurred, we observed that fish in **Group 2** (water supplemented with W1) had significantly higher weight, length, and height compared to the control group, while the other experimental groups did not significantly differ from the control. The literature shows mixed results regarding the effect of probiotic compounds on the growth performance of carp. In a 31-day feeding experiment Tang et al. (2016) did not observe a significant effect of feed supplementation with commercial supplements on carp growth performance. In a field experiment reported by Mohammadian et al. (2022), no significantly better carp growth performance was observed under an experimental diet supplemented with *Lactobacillus* species. On the other hand, the positive effects of targeted supplementation with specific genera have been reported. Supplementation with *Lactococcus* improved the growth performance of common carp (Feng et al., 2019, 2022). A positive effect of *Lactobacillus* supplementation was reported by Soltani et al. (2017), Yassine et al. (2021), and Zhang et al. (2019). Also experimental diets with selected *Bacillus* species had a positive impact on carp’s growth Yanbo and Zirong, 2006). A positive effect on growth based on feeding commercial probiotic supplements was reported by Ghasempour Dehaghani et al. (2015). These authors suggested that the effect is due to a positive influence of the supplementation on glucose absorption and overall metabolism of the carp, as well as due to better health status due to improved immune resistance of the fish.

The contradiction in the literature reports can be attributed to two main reasons. First, applying experimentally prepared diets that contained a specific composition of species in a predefined concentration may substantially differ from supplementation with commercially available products for which the microbial composition is often not well defined and not specifically controlled in each particular sample. This problem was demonstrated in our study, where sequencing results of commercial supplements did not detect all the microorganisms that were listed as the designated components, indicating that some of the genera might be present in a very low abundance under the technical detection threshold. The second aspect is that most of the above-mentioned studies (except Mohammadian et al. (2022)) were carried out in experimental conditions, in tanks. Especially in the context of fish growth it is expected that such environmentally unified conditions strongly deviate from the situation in the field. Although we are not aware of the systematic comparison regarding tank and experimental growth performance of fish, due to environment-microbiome-phenotype interactions that are not fully modelled in tank based experiments, experimental results may not be directly transferable to the field.

Given our observation of no impact of the supplementation with effective microorganisms on overall bacterial community composition, we also estimated the impact of particular families on fish growth performance. The highest significant positive effect on weight with the increased abundance of *Flavobacteriales* may be related to the fact that members of this typically highly abundant family can “assist” probiotic supplementation by degrading polysaccharides (Shan et al., 2025). Conversely the highest negative effect on fish weight and length with the increased abundance was estimated for *Silvanigrellaceae*, that has not been characterised in the context of aquaculture. Somewhat surprisingly, a positive effect on weight and length was estimated for the increased abundance of the *Nocardiaceae* family. Surprisingly, some Nocardia species cause nocardiosis, a serious disease in aquaculture. However this family also contains species that play a role in organic matter turnover and in the degradation of xenobiotics that may provide beneficial effects for the fish (summarised in Rosenberg et al., 2014d).

## 5. Conclusion

Although the effective microorganisms supplementation with a commercial product did not induce significant modifications in water, sediment or gut microbiome of the fish, still an increase in the final weight was observed in the fish reared the ponds supplemented with W1. This experimental group (**Group 2**) was distinct from the other experimental conditions because it showed the lowest and most consistent alpha diversity of water and gut across ponds, and appeared to be most stable in diversity across the course of the experiment. We hypothesise that the supplementation of water provided an improved microbial environment that also remained stable throughout the course of the experiment – what appeared to be beneficial for the fish.

## Acknowledgements

The study was supported by the National Science Centre, Poland, under the research project 2021/41/B/NZ9/01409. Sequencing was performed thanks to Genomics Core Facility CeNT UW (RRID:SCR_022718), using NovaSeq 6000 platform financed by Polish Ministry of Science and Higher Education (decision no. 6817/IA/SP/2018 of 2018-04-10). We thank the developers of artificial intelligence tools that assisted in data visualisation for this study.

## Data Availability

The sequence reads will be available from the NCBI BioProject database under the accession ID xxx upon acceptance of the manuscript.

## Conflict of interest

The authors declare no conflict of interests.

## Authorship contribution statement

**Michalina Jakimowicz:** Software, Formal analysis, Data Curation, Writing – review and editing. **Katarzyna Sidorczuk:** Software, Formal analysis, Data Curation, Writing – review and editing. **David Huyben:** Conceptualization, Writing – review and editing. **Falk Hildebrand:** Software, Methodology, Writing – original draft, Writing – review and editing. **Łukasz Napora–Rutkowski:** Conceptualization, Data Curation, Methodology, Project administration, Resources, Writing – review and editing. **Piotr Hajduk:** Data curation, Formal analysis. **Marek Sztuka:** Data curation, Writing – original draft. **Magda Mielczarek:** Supervision, Writing – original draft. **Dawid Słomian:** Data curation, Formal analysis. **Urszula Szulc:** Data curation, Formal analysis. **Laura Jarosz:** Data curation, Formal analysis. **Joanna Szyda:** Conceptualization, Methodology, Supervision, Project administration, Visualization, Writing - Original Draft, Writing – review and editing.

## Funding

This research and the APC were funded by The National Science Centre (NCN), grant number 2021/41/B/NZ9/01409. KS and FH were supported by the European Research Council H2020 StG (erc-stg-948219, EPYC), the Quadram Institute Bioscience’s Institute Strategic Programme (ISP) BB/X011054/1, workpackage BBS/E/F/000PR13631 and the Earlham ISP BBX011089/1, projects BBS/E/ER/230002A and BBS/E/ER/230002B.

## Notes

### Competing Interest Statement

Michalina Jakimowicz, Piotr Hajduk, Marek Sztuka, Magda Mielczarek, Dawid Słomian, and Joanna Szyda received financial support from the National Science Centre, Poland.
All other authors declare no competing interests.

